# Distinct functions of cardiac β-adrenergic receptors in the T-tubule *vs.* outer surface membrane

**DOI:** 10.1101/2021.04.28.441732

**Authors:** Marion Barthé, Flora Lefebvre, Emilie Langlois, Florence Lefebvre, Patrick Lechêne, Xavier Iturrioz, Catherine Llorens-Cortes, Tâp Ha-Duong, Laurence Moine, Nicolas Tsapis, Rodolphe Fischmeister

**Author notes:** Corresponding author: Rodolphe Fischmeister, PhD, **E-mail:**. **Author contributions:** Conceptualisation, R.F. and N.T.; Methodology, M.B., N.T., R.F., L.M., T.H.D. and X.I.; Validation, R.F., L.M. and N.T.; Formal Analysis, M.B., F.L. and E.L.; Investigation, M.B., F.L., E.L., F.L., P.L., X.I., T.H.D. and L.M.; Resources, M.B., F.L., E.L. and T.H.D.; Writing, M.B., N.T., L.M., T.H.D and R.F.; Visualisation, M.B.; Supervision N.T. and R.F.; Project Administration N.T. and R.F.; Funding Acquisition N.T. and R.F.

## Abstract

β-adrenoceptors (β-ARs) regulate cardiac function during sympathetic nerve stimulation. β-ARs are present in both cardiac T-tubule (TTM) and outer surface membrane (OSM), but how their location impacts on their function is unknown. Here, we developed a technology based on size exclusion to explore the function of β-ARs located in the OSM. We synthetized a PEG-Iso molecule by covalent linking isoprenaline (Iso) to a 5000 Da PolyEthylene-Glycol (PEG) chain to increase the size of the β-AR agonist and prevent it from accessing the TT network. The affinity of PEG-Iso and Iso on β_1_- and β_2_-ARs was measured using radioligand binding. Molecular dynamics simulation was used to assess PEG-Iso conformation and visualise the accessibility of the Iso moiety to water. Using confocal microscopy, we show that PEGylation constrains molecules outside the T-tubule network due to the presence of the extracellular matrix. β-AR activation in OSM with PEG-Iso produced a lower stimulation of [cAMP]_i_ than Iso but a larger stimulation of cytosolic PKA at equivalent levels of [cAMP]_I_ and similar effects on excitation-contraction coupling parameters. However, PEG-Iso produced a much lower stimulation of nuclear PKA than Iso. Thus, OSM β-ARs control mainly cytosolic cAMP/PKA pathway and contractility, while TTM β-ARs control mainly nuclear PKA and nuclear protein phosphorylation. Size exclusion strategy using ligand PEGylation provides a unique approach to evaluate the respective contribution of T-tubule *vs.* outer surface membrane proteins in cardiac cells.

**Significance Statement:** β-adrenoceptors (β-ARs) regulate cardiac function during sympathetic nerve stimulation. They are present in both cardiac T-tubule and outer surface membranes, but how their location impacts on their function is unknown. By linking the β-AR agonist isoprenaline (Iso) to a PolyEthylene-Glycol (PEG) chain, we increased the size of the agonist to prevent it from entering the T-tubules. Thus, PEG-Iso is only able to activate β-ARs in the outer surface membrane. With this size exclusion strategy, we show that β-ARs located in the outer surface membrane control mainly cytosolic cAMP/PKA pathway and contractility, while those located in the T-tubule membrane control mainly nuclear PKA and nuclear protein phosphorylation.

## introduction

The β-adrenergic receptor (β-AR) is a key player in the regulation of cardiac function during sympathetic nerve stimulation. The classical pathway for β-AR receptor signalling is activation of adenylyl cyclases (AC) via G_αs_, resulting in increased intracellular cAMP levels ([cAMP]_i_) (46). Three types of β-ARs are expressed in the heart, respectively β_1_-, β_2_- and β_3_-ARs. β_1_- and β_2_-ARs are both positively coupled to AC/cAMP cascade and cardiac performance (76) while β_3_-ARs may act via either cAMP of cGMP depending on the cell type (atrial *vs.* ventricular myocytes) (61, 64). The primary target of cAMP is the cAMP-dependent protein kinase (PKA) that in turn phosphorylates several key proteins involved in the excitation-contraction (EC) coupling, such as the L-type Ca^2+^ channel (LTCC or Ca_V_1.2), phospholamban, troponin I, etc. (9). The phosphorylation of Ca_v_1.2 (or its regulatory protein Rad (45, 47)) leads to enhanced LTCC current (*I*_Ca,L_) and sarcoplasmic reticulum (SR) Ca^2+^ release via the ryanodine type 2 receptor (RyR2), contributing to enhanced Ca^2+^ transients and contraction (8, 24). Phosphorylation of phospholamban increases Ca^2+^ uptake into the SR which accelerates Ca^2+^ transient decay and, together with troponin I phosphorylation, speeds up relaxation. Whereas short term stimulation of β-AR/cAMP is beneficial for the heart, chronic activation of this pathway results in altered Ca^2+^ signalling, cardiac hypertrophy and fibrosis, leading to ventricular dysfunction (55) and cardiac arrhythmias (4, 11, 25, 40, 51).

The cell membrane of cardiomyocytes is characterized by invaginations of the surface membrane, occurring primarily perpendicular to myocyte longitudinal edges, at intervals of ~1.8–2 µm, that form a complex interconnected tubular network penetrating deep into the cell interior (36). This network is called the transverse (T−) tubules (TT) system, although the invaginations may often bifurcate in the axial direction or form branches (36). These tubular structures are found mostly in adult ventricular myocytes, where they represent about 30% of the total cell membrane (12, 17), and occur near the sarcomeric z-discs (18), allowing functional junction with the SR called dyad. They are critical for EC coupling by concentrating LTCCs and positioning them at close proximity of RyR2 clusters at the junction of SR to form Ca^2+^ release units (31, 63). During an action potential, TT propagate the cell-membrane depolarization inside the cell allowing Ca^2+^ entry to trigger successive Ca^2+^ release thus promoting synchronicity as well as efficiency of the Ca^2+^-induced Ca^2+^ release (CICR) phenomenon (28). Their enrichment with ion channels and the presence of signaling pathways components such as β-ARs (53) and ACs (71) make these structures essential for cardiomyocyte function and regulation.

Biochemical assays following membrane fractionation have provided indirect evidence that membrane proteins may have different properties whether located in TTM or OSM. Other indirect evidence came from experiments in cardiomyocytes in which TTM was uncoupled from OSM (detubulation) using a hyperosmotic shock with molar concentrations of formamide (13, 14, 16, 17, 20, 21, 27, 41, 50, 70). However, none of these approaches allowed to investigate in an intact cardiomyocyte whether the function of a given membrane receptor differs if the receptor is located in OSM or TTM. To do so requires to being able to separately activate or inhibit the receptor in OSM or TTM which has not been feasible so far. Here, we tackled this challenge by developing a size exclusion strategy using the PEGYlation technology. By a covalent link between isoprenaline (Iso) and a PolyEthylene-Glycol (PEG) chain, we increased the size of the β-AR agonist to prevent it from accessing the TT network. Our working hypothesis is that PEGylated isoprenaline (PEG-Iso) activates only β-ARs present in OSM while free Iso activates β-ARs present in both OSM and TTM. We thus characterized the properties of PEG-Iso and compared its functional effects in adult rat ventricular myocytes (ARVMs) with those of Iso.

## Results

### PEGylation constrains molecules outside the TT network

The rationale for using ligand PEGylation as a tool to impede ligand diffusion into T-tubules came from the observation that fluorescent PEG_5000_ molecules were unable to access to TTs. A typical experiment (out of 15) using confocal microscopy is shown in Fig. 1. When ARVMs were exposed during 15 min to 100 µM of PEG_5000_ molecules functionalized with Fluorescein Isothiocyanate (FITC), no fluorescence was seen throughout the cell (Fig. 1A, B). On the contrary, when the cells were exposed during 15 min to 100 µM fluorescein, fluorescence was seen with clear staining of the TTs (Fig. 1C, D). We suspected that the lack of access of PEG-FITC in the TTs was due to steric hindrance due to the presence of glycocalyx matrix. To test this hypothesis, ARVMs were first exposed during 1 h to a solution containing 0.25 U/mL neuraminidase, an enzyme that degrades sialic acid in the glycocalyx matrix (56). As shown in Fig. 1E, F, this allowed PEG-FITC to enter the TTs as evidenced by the striated profile now observed.

**Fig. 1.**
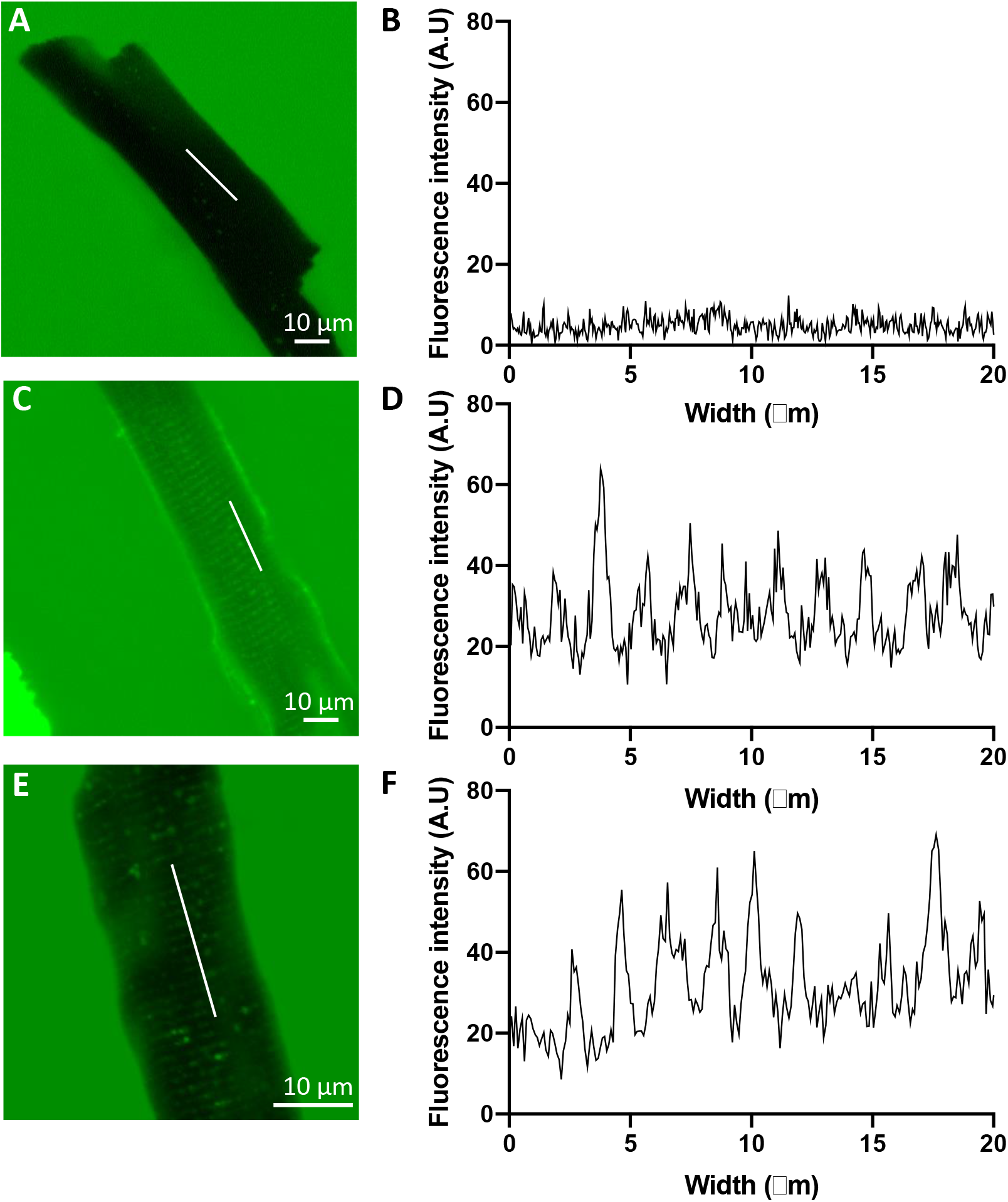
Localization of PEG-FITC in freshly isolated adult rat ventricular cardiomyocytes. Typical confocal images (**A**, **C**, **E**) and plot profiles representing the fluorescence intensity measured across the cell (**B**, **D**, **F**) of ARVMs incubated during 15 min with either 100 µM PEG_5000_-FITC (**A, B**), 100 µM free fluorescein (**C**, **D**), or 100 µM PEG_5000_-FITC after a 1h treatment with 0.25 U/mL neuraminidase (**E**, **F**).

### PEG-Isoprenaline binds to β_1_- and β_2_-adrenergic receptors

The above experiments demonstrate that PEG_5000_ molecules are unable to access the TTs in our experimental conditions. We thus synthetized PEGylated isoprenaline (PEG-Iso) by linking isoprenaline to PEG_5000_ molecules (Fig. S1). Our rationale was that since PEG-Iso does not access TTs, it will not reach β-ARs located in the TTM. But the question remained whether PEG-Iso would be able to bind to β-ARs located in the OSM. To address this question, we performed radioligand binding studies in purified membranes from Chinese Hamster Ovary (CHO) cells overexpressing either β_1_- or β_2_-ARs. Competition curves between [^125^Iodo]cyanopindolol and Iso or PEG-Iso were used to measure K_i_ values for both ligands. As shown in Fig. 2A and B, PEG-Iso binds to both β_1_- and β_2_-ARs, but with an affinity which is ~2-orders of magnitude lower than Iso. We hypothesized that the decreased affinity of PEG-Iso could be due to wrapping of the PEG chain around Iso moiety reducing its exposure to solvent (water). To test this, we used molecular dynamics simulations to evaluate the solvent-accessible surface area (SASA) of Iso moiety in PEG-Iso compared to free Iso. Fig. 2C shows that free Iso SASA remains steady over the whole studied time range (black line). In contrast, Iso moiety in PEG-Iso (blue line) has a SASA which largely fluctuates and reaches many times values close to zero, indicating its frequent shielding by the PEG chain. We calculated that only 1.2% of PEG-Iso conformations have an Iso SASA larger than 90% of the free Iso average value. This means that over a time lapse of 100 s, the Iso moiety is unwrapped from the PEG chain and therefore able to bind to a receptor during only a short time of about 1 s.

**Fig. 2.**
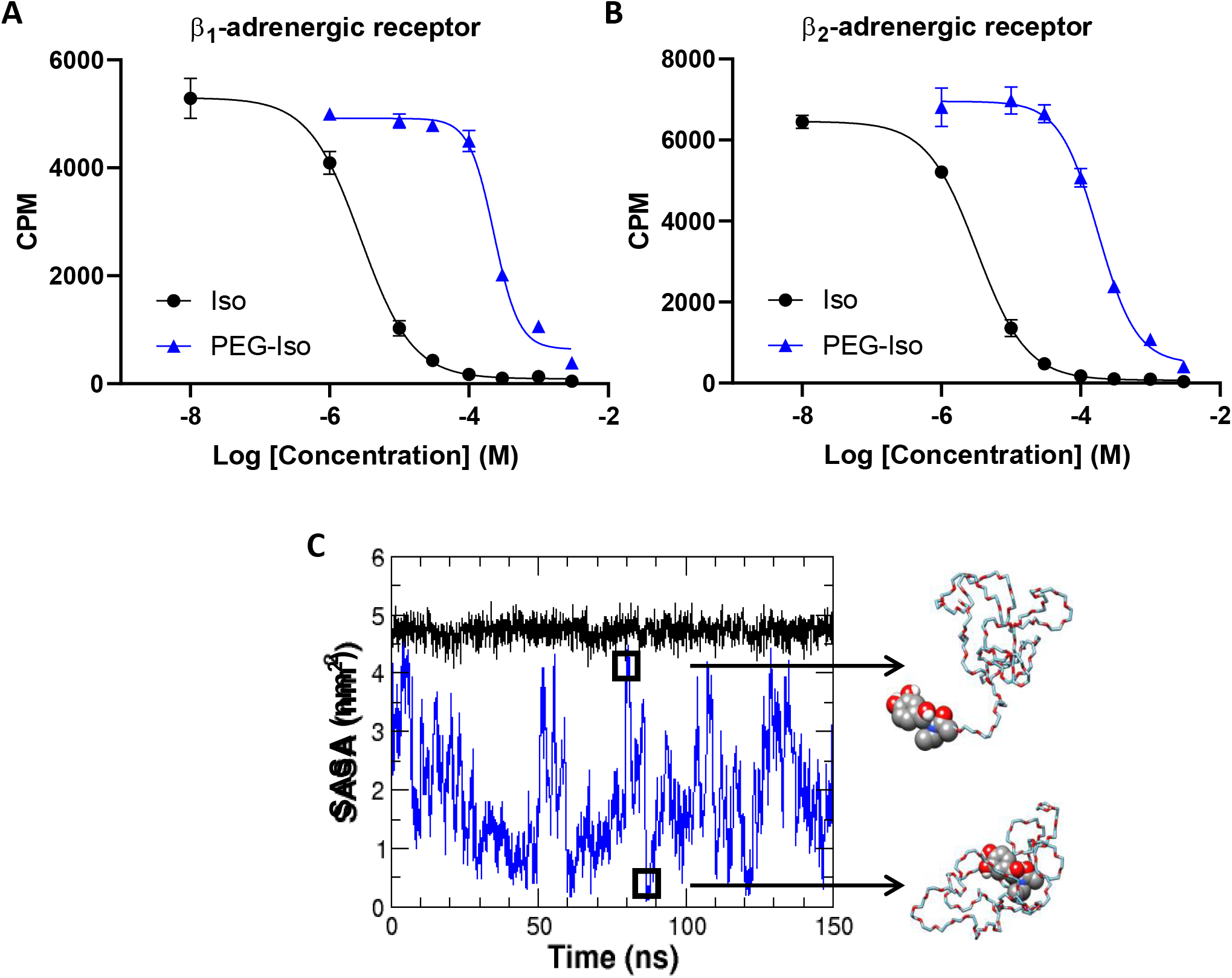
Binding properties of Iso and PEG-Iso on ß-ARs. ß_1_- or ß_2_-AR CHO membrane preparations (2.5 μg/triplicate) were incubated for 2h with 0.25 nM [^125^Iodo]cyanopindolol and increasing concentrations of unlabeled Iso or PEG-Iso. (**A**) Example of a typical competition binding of Iso and PEG-Iso on ß_1_-ARs. (**B**) Example of a typical competition binding of Iso and PEG-Iso on ß_2_-ARs. K_i_ values were, respectively, 2.0 ± 0.7 µM and 1.4 ± 0.7 mM for Iso and PEG-Iso on ß_1_-ARs; 2.1 ± 0.4 µM and 429 ± 1 µM for Iso and PEG-Iso on ß_2_-ARs (N=5). (**C**) Time evolution of solvent-accessible surface area (SASA) of Iso (black trace) and PEG-Iso (blue trace) using molecular dynamics simulations. Two representative conformations of PEG-Iso with large or small SASA are shown.

### Comparison of the effects of PEG-Iso and Iso on cytosolic cAMP

The next series of experiments were designed to test whether PEG-Iso was able to produce a functional β-AR response in ARVMs. First, cytosolic cAMP ([cAMP]_i_) was monitored in isolated ARVMs expressing the FRET-based sensor Epac-S^H187^ (42). As seen in Fig. 3, both Iso (Fig. 3A, B) and PEG-Iso (Fig. 3C, D) produced a concentration-dependent increase in [cAMP]_i_. However, there were two major differences: 1) the concentration-response curve to PEG-Iso was shifted ~100-fold towards larger concentrations as compared to Iso; 2) the maximal efficacy of PEG-Iso was significantly lower by ~30% than that of Iso. The former was anticipated based on the lower binding affinity of PEG-Iso to β-ARs shown above. The latter can be explained by the fact that PEG-Iso only activates β-ARs in OSM while Iso activates β-ARs in both OSM and TTM, thus producing a larger [cAMP]_i_ elevation.

**Fig. 3.**
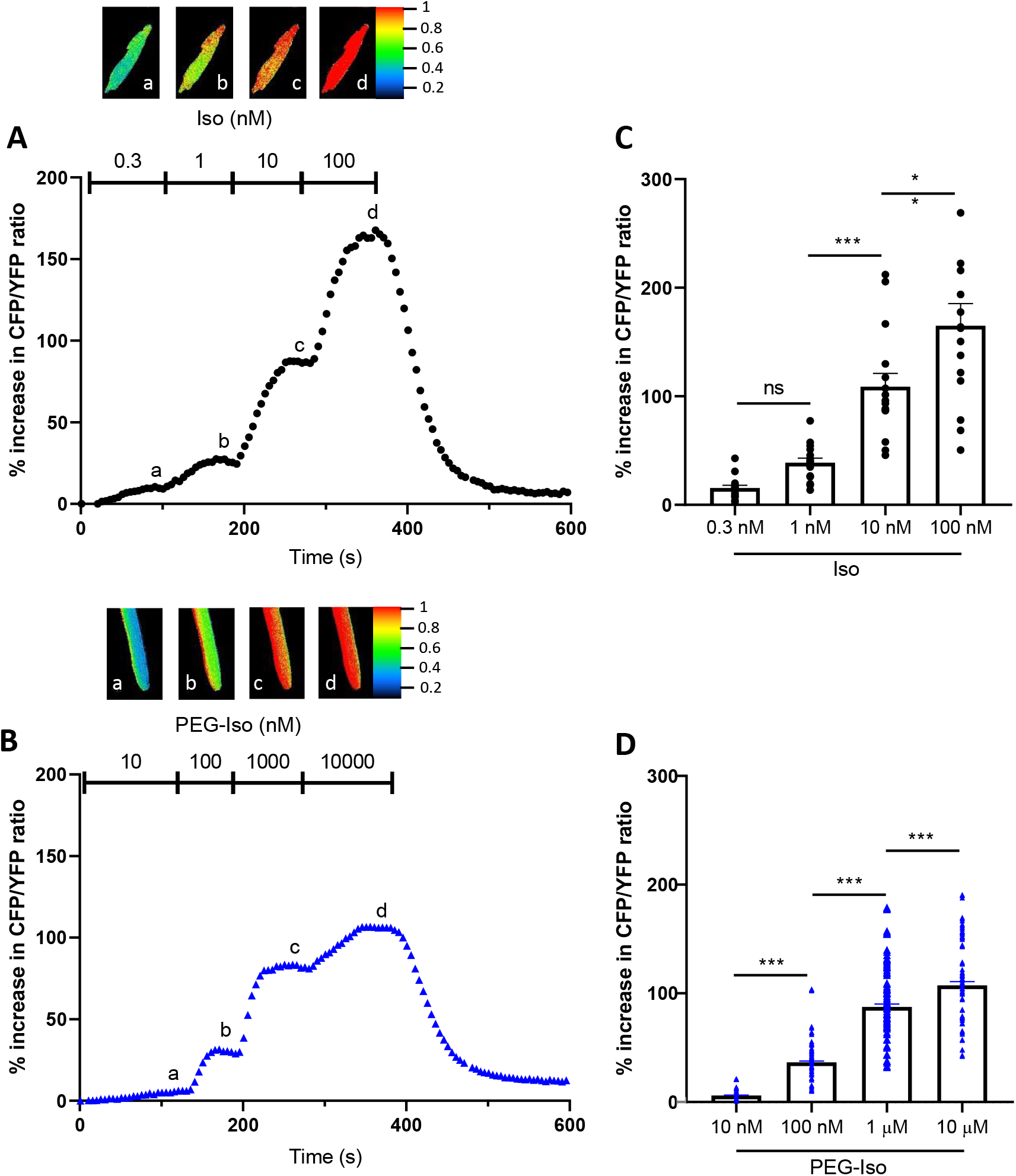
Effect of Iso and PEG-Iso on cytosolic cAMP in adult rat ventricular cardiomyocytes. Freshly isolated ARVMs were infected with an adenovirus encoding the Epac-S^H187^ FRET-based cytosolic cAMP sensor for 48h at 37°C at a multiplicity of infection of 1000 pfu/cell. (**A**) Typical experiment showing the time course of CFP/YFP ratio during successive applications of four increasing Iso concentrations: 0.3, 1, 10 and 100 nM. (**B**) Similar experiment showing the time course of CFP/YFP ratio during successive applications of four increasing PEG-Iso concentrations: 10 and 100 nM, 1 and 10 µM. Pseudo-color images shown above the main graphs were taken at times indicated by the corresponding letters in the graphs. (**C**, **D**) Summary data from several similar experiments as in (**A**) and (**B**), respectively. The bars show the mean ± s.e.m of the data shown by symbols. 16 cells from 3 rats were used in (**C**); 32 cells from 4 rats in (**D**). One-way ANOVA and Tukey’s multiple comparisons post hoc test were used: ** p<0.01; *** p<0.001; ns, non significant.

### Comparison of the effects of PEG-Iso and Iso on *I*_Ca,L_

Next, the ability of Iso and PEG-Iso to stimulate the L-type Ca^2+^ current, *I*_Ca,L_ was compared at a single concentration of each, i.e. 10 nM Iso and 1 µM PEG-Iso, shown above to produce an equivalent response on [cAMP]_i_ (Fig. 4). As shown in the individual experiments in Fig. 4A and B, both Iso and PEG-Iso produced an increase in the amplitude of *I*_Ca,L_, and both responses were on average similar in amplitude (Fig. 4C).

**Fig. 4.**
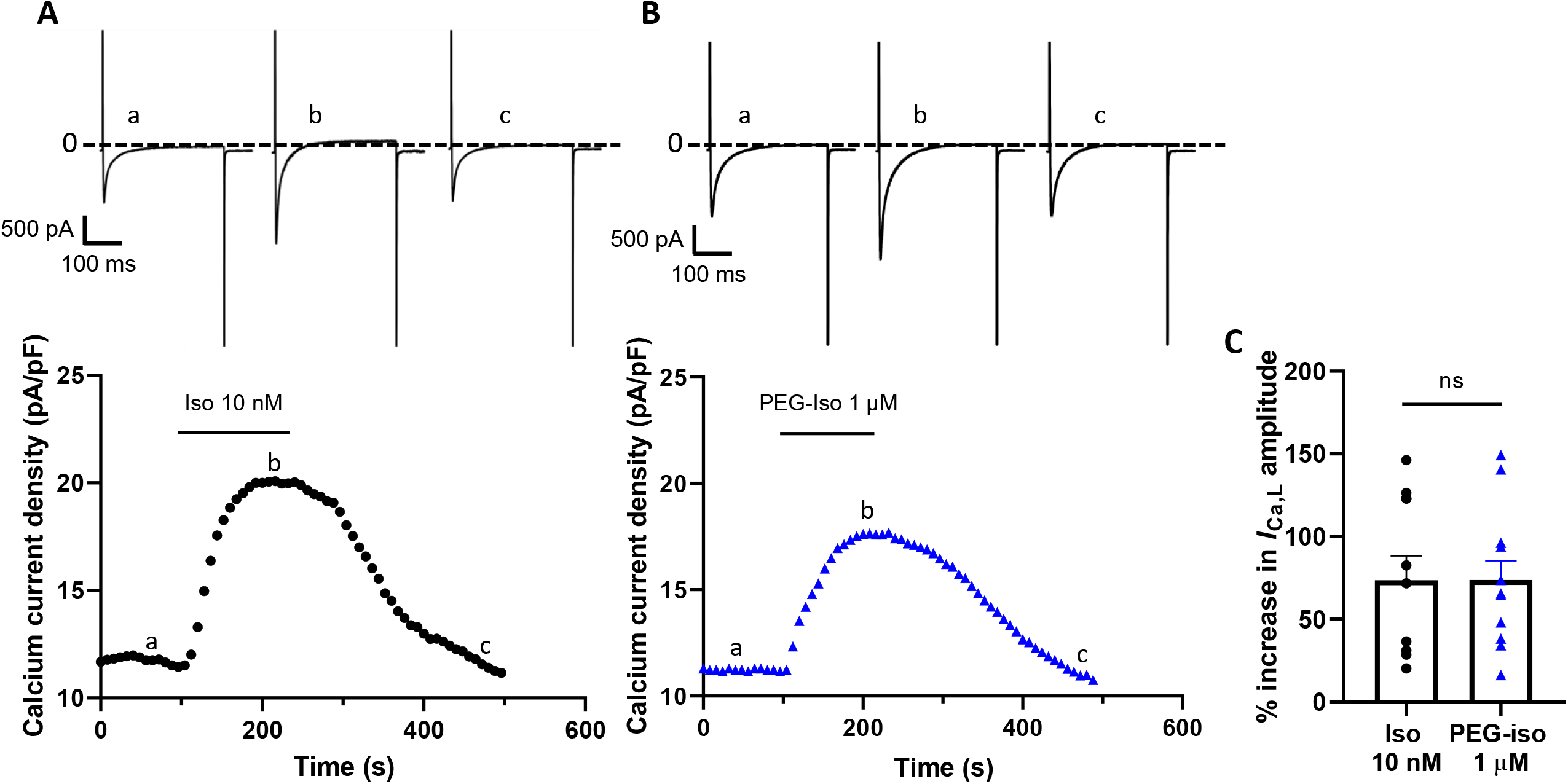
Comparison of the effects of PEG-Iso and Iso on *I*_Ca,L_. The whole-cell patch-clamp technique was applied to ARVMs after 24 h culture. (**A**) Typical experiment showing the time course of *I*_Ca,L_ current density upon application of 10 nM Iso and during washout. (**B**) Similar experiment showing the response of *I*_Ca,L_ current density to 1 µM PEG-Iso. Individual current traces shown above the graphs were taken at times indicated by the corresponding letters on the graphs. **c** Summary data from several similar experiments as in (**A**) and (**B**). Student *t*-test: ns, non significant.

### Comparison of the effects of PEG-Iso and Iso on excitation-contraction coupling

To investigate the impact of a stimulation of OSM β-ARs on EC coupling, Ca^2+^ transients and sarcomere shortening were simultaneously recorded in Fura-2-loaded ARVMs and their response to PEG-Iso and Iso were evaluated. Here, two concentrations of PEG-Iso were used, 100 and 300 nM, and two concentrations of Iso producing similar elevations of [cAMP]_i_, 1 and 3 nM, respectively. As shown in Fig. 5 and 6, 100 and 300 nM PEG-Iso increased contractility (Fig. 5B), Ca^2+^ transients (Fig. 6B) and accelerated their relaxation kinetics (Fig. 5D and 6D). These effects were not different from those produced by 1 and 3 nM Iso, respectively, on either sarcomere shortening (Fig. 5A, C), Ca^2+^ transient amplitude (Fig. 6A, C) or relaxation kinetics (Fig. 5C, D; Fig. 6C, D).

**Fig. 5.**
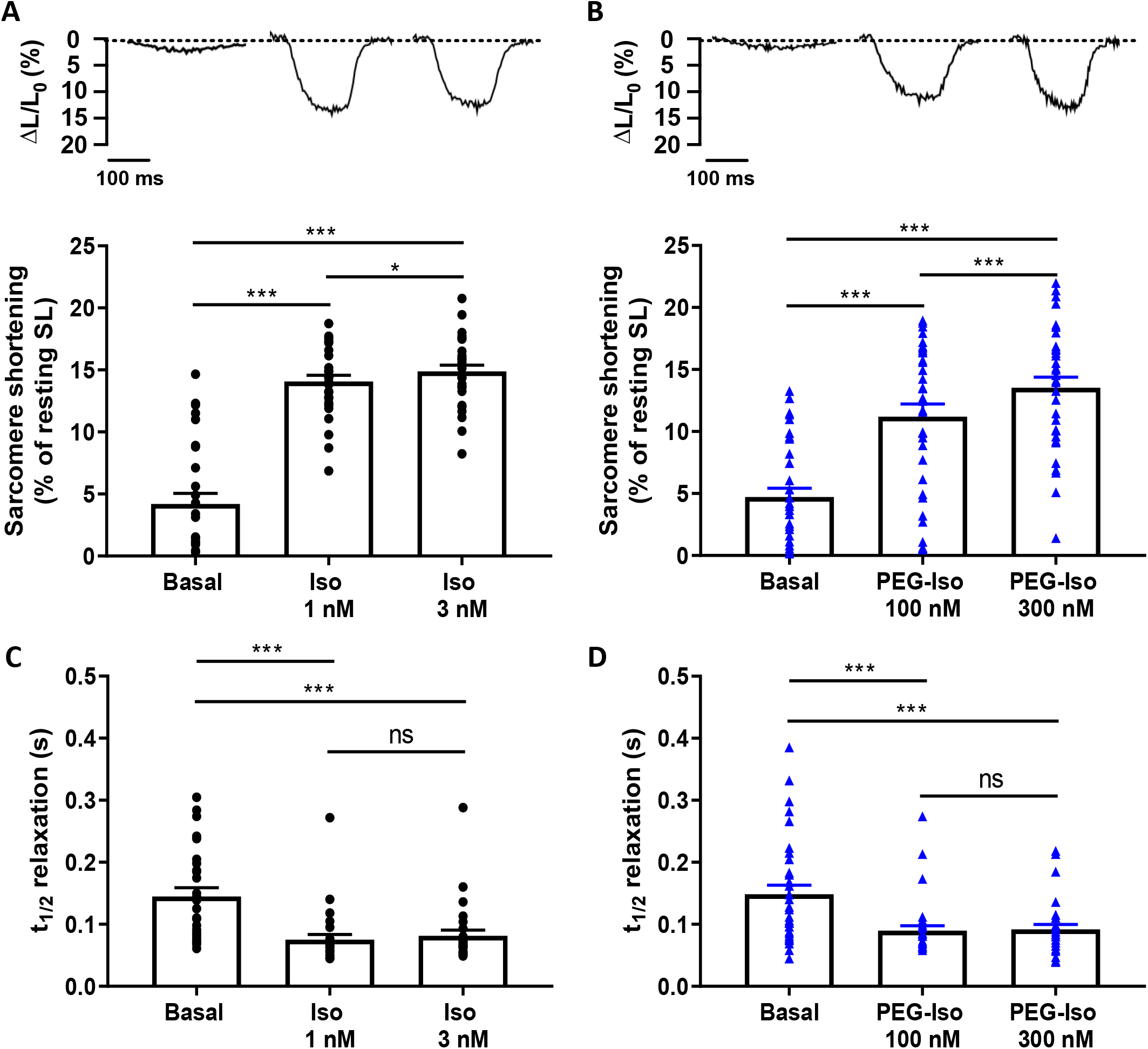
Comparison of the effects of PEG-Iso and Iso on sarcomere shortening. Representative traces of sarcomere shortening recorded in ARVMs paced at 0.5 Hz and loaded with Fura-2 AM (1 µM) showing the effect of Iso (1 and 3 nM, **A**) and PEG-Iso (100 and 300 nM, **B**). The bars in (**A**) and (**B**) show the mean ± s.e.m of the data shown by symbols. (**C**, **D**) Average time-to-50% relaxation of sarcomere shortening from experiments shown in (**A**) and (**B**), respectively. 30 cells from 4 rats were used in (**A**) and (**C**); 34 cells from 4 rats in (**B**) and (**D**). One-way ANOVA and Tukey’s post hoc test: * p<0.05; *** p<0.001; ns, non-significant.

**Fig. 6.**
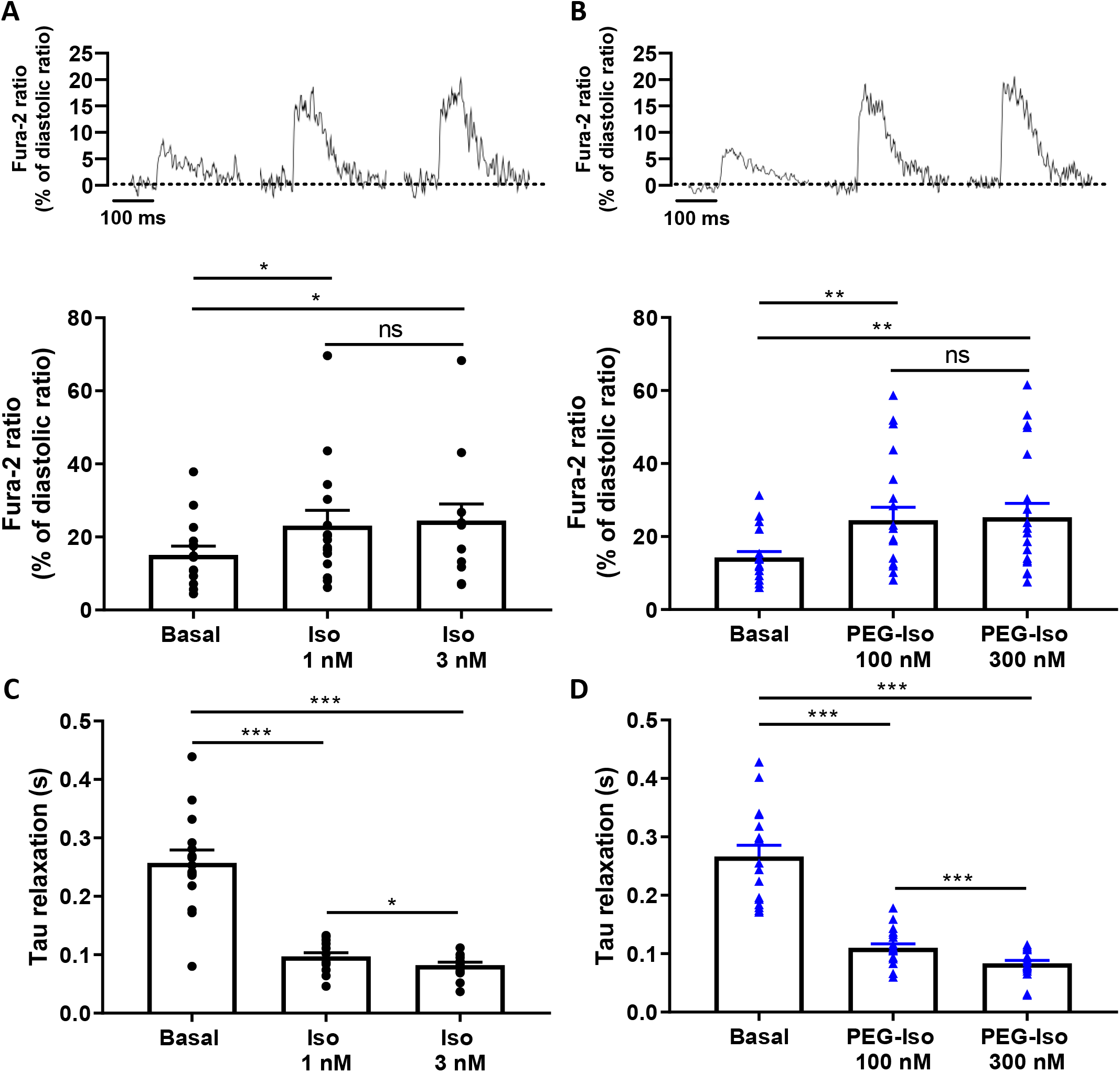
Comparison of the effects of PEG-Iso and Iso on Ca^2+^ transients. Representative traces of Ca^2+^ transients recorded in ARVMs paced at 0.5 Hz and loaded with Fura-2 AM (1 µM) showing the effect of Iso (1 and 3 nM, **A**) and PEG-Iso (100 and 300 nM, **B**). The bars in (**A**) and (**B**) show the mean ± s.e.m of the data shown by symbols. (**C**, **D**) Exponential time constant (*Tau*) of relaxation of Ca^2+^ transients from experiments shown in (**A**) and (**B**), respectively. 30 cells from 4 rats were used in (**A**) and (**C**); 34 cells from 4 rats in (**B**) and (**D**). One-way ANOVA and Tukey’s post hoc test: * p<0.05; ** p<0.01; *** p<0.001; ns, non-significant.

### Comparison of the effects of PEG-Iso and Iso on cytosolic and nuclear PKA activity

The observation that equipotent concentrations of PEG-Iso and Iso on [cAMP]_i_ produced equivalent effects on *I*_Ca,L_ and EC coupling, may suggest that OSM β-ARs are not functionally different from their TTM homologs as long as their degree of activation leads to similar [cAMP]_i_ responses. However, the above experiments do not exclude possible differences in the compartmentalization of intracellular cAMP cascade when cAMP is synthesized upon OSM or TTM β-AR stimulation. To obtain further insight, we compared the effect of PEG-Iso and Iso on PKA activity in the bulk cytoplasm and in the nucleus. For that, we used genetically encoded A-kinase activity FRET-based reporters (AKAR3) targeted to these compartments by the addition of a nuclear export sequence (NES), and a nuclear localizing sequence (NLS), respectively (3). As shown previously (6, 32), adenoviral transfer allowed robust and compartment-specific expression of these biosensors after 24 h in ARVMs. In the experiments shown in Fig. 7, four increasing concentrations of Iso (0.3, 1, 3 and 10 nM: Fig. 7A, C) and PEG-Iso (10, 30, 100 nM and 1 µM: Fig. 7B, D) were successively applied to ARVMs infected with AKAR3-NES probe and cytosolic PKA activation was measured. Both compounds produced a clear concentration-dependent increase in cytosolic PKA activity, with a similar efficacy but a ~100-fold lower potency for the effect of PEG-Iso as compared to Iso. However, when nuclear PKA activity was measured, a clear difference between the effects of Iso and PEG-Iso were observed (Fig. 8): While PEG-Iso and Iso still produced a concentration-dependent increase in nuclear PKA activity, the response to 1 µM PEG-Iso (Fig. 8D) was reduced about 2-fold as compared to 10 nM Iso (Fig. 8C). The same difference is observed regarding the nuclear proteins phosphorylation levels (Fig. S2) after stimulation with 10 nM Iso or 1 µM PEG-Iso.

**Fig. 7.**
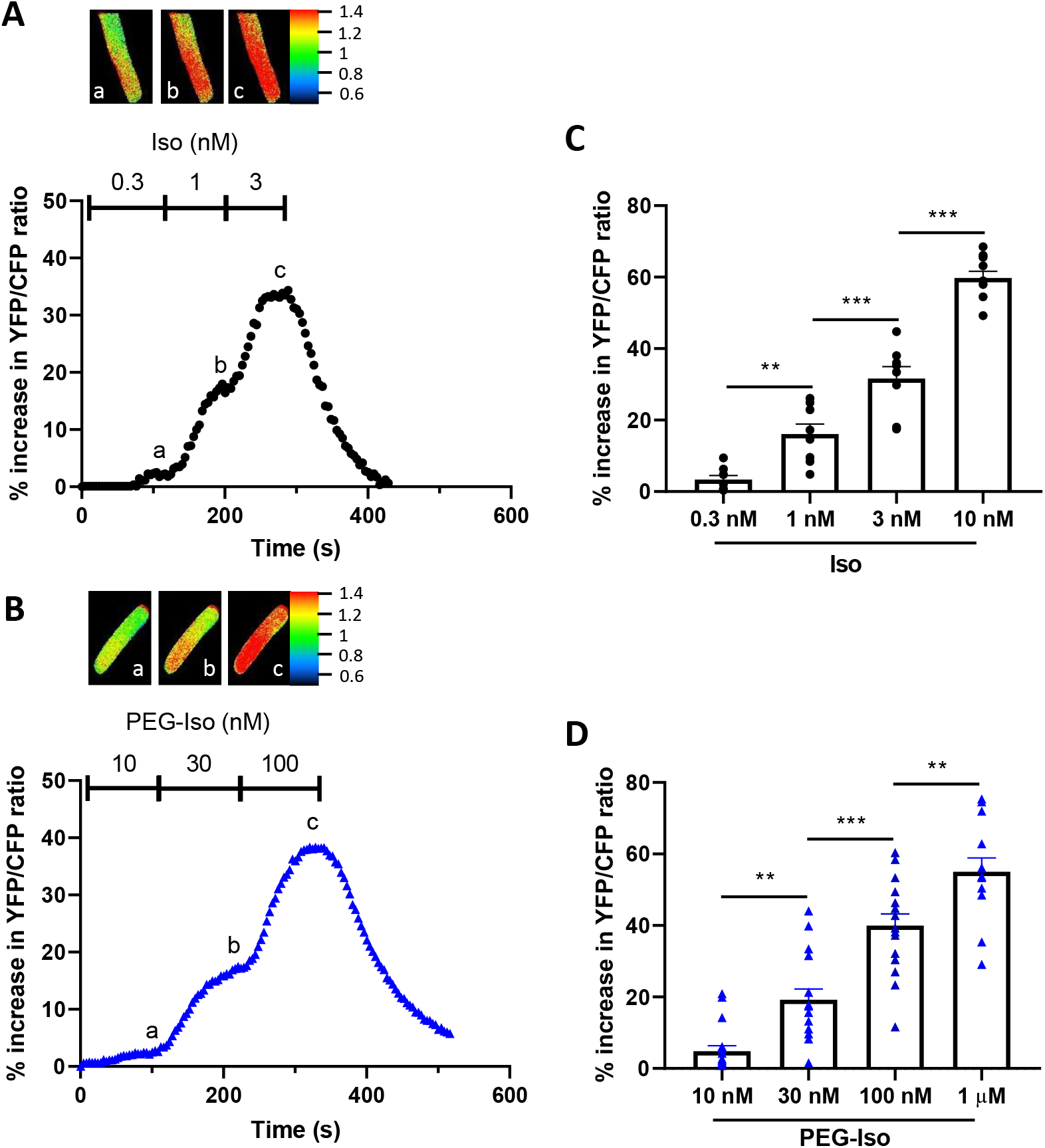
Comparison of the effects of PEG-Iso and Iso on cytosolic PKA activity. Freshly isolated ARVMs were infected with an adenovirus encoding the AKAR3-NES FRET-based PKA sensor for 48h at 37°C at a multiplicity of infection of 1000 pfu/cell. (**A)** Typical experiment showing the time course of YFP/CFP ratio during successive applications of three increasing Iso concentrations: 0.3, 1 and 3 nM. (**B**) Similar experiment showing the time course of YFP/CFP ratio during successive applications of three increasing PEG-Iso concentrations: 10, 30 and 100 nM. Pseudo-color images shown above the main graphs were taken at times indicated by the corresponding letters in the graphs. (**C**, **D**) Summary data from several similar experiments as in (**A**) and (**B**), respectively. The bars show the mean ± s.e.m of the data shown by symbols. 3 rats and 8-10 cells were used in **c**; 3-5 rats and 13-18 cells in (**D**). One-way ANOVA and Tukey’s multiple comparisons post hoc test: ** p<0.01; *** p<0.001.

**Fig. 8.**
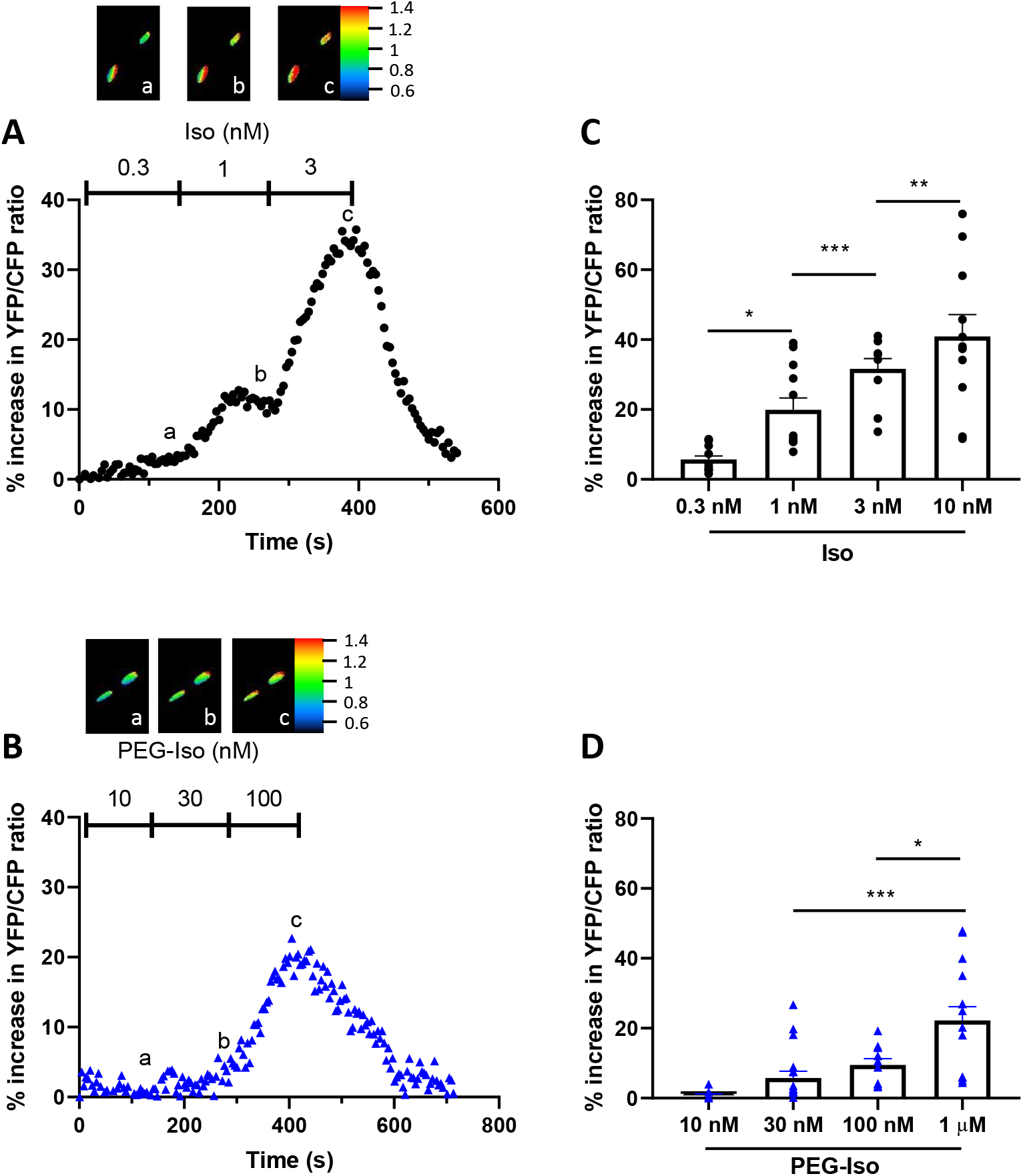
Comparison of the effects of PEG-Iso and Iso on nuclear PKA activity. Freshly isolated ARVMs were infected with an adenovirus encoding the AKAR3-NLS FRET-based PKA sensor for 48h at 37°C at a multiplicity of infection of 1000 pfu/cell. (**A**) Typical experiment showing the time course of YFP/CFP ratio during successive applications of three increasing Iso concentrations: 0.3, 1 and 3 nM. (**B**) Similar experiment showing the time course of YFP/CFP ratio during successive applications of three increasing PEG-Iso concentrations: 10, 30 and 100 nM. Pseudo-color images shown above the main graphs were taken at times indicated by the corresponding letters in the graphs. (**C**, **D**) Summary data from several similar experiments as in (**A**) and (**B**), respectively. The bars show the mean ± s.e.m of the data shown by symbols. 3 rats and 10-12 cells were used in **c**; 3-5 rats and 10-17 cells in (**D**). One-way ANOVA and Tukey’s multiple comparisons post hoc test: * p<0.05; ** p<0.01; *** p<0.001.

Fig. S3 shows a plot of cytosolic and nuclear PKA activity as a function of cytosolic cAMP from the average data in the experiments in Fig. 3, 7 and 8 for 3 concentrations of PEG-Iso (10 and 100 nM and 1 µM) and Iso (0.3, 1 and 10 nM). It shows that for any given measured increase in [cAMP]_i_, PEG-Iso is more efficient than Iso to increase PKA activity in the cytosol (Fig. S3A), while on the contrary Iso is more efficient than PEG-Iso to increase PKA activity in the nucleus (Fig. S3B).

## Discussion

The main function of the TT system is to provide proximity between LTCCs in the TTM and RyR2 in the SR membrane (19, 63). However, TTM also contains many other membrane proteins, such as receptors, enzymes, ion channels and transporters. These include plasma membrane Ca^2+^-ATPase (15), Na^+^-Ca^2+^ exchanger (62, 77), Na^+^ channels (62), Na^+^/K^+^-ATPase (7), ACs (29, 71) and β-ARs (30, 53). Density of these membrane proteins is usually found to be higher in TTM than in the external membrane. Hence, the TT network plays a determinant role in rapid activation and synchronous Ca^2+^ release, cellular signalling and, consequently, cardiac contraction in both human and animal models (34, 67, 68).

Any given membrane protein is likely to serve different functions and be regulated in a different manner whether it is located in TTM or in OSM. Attempts to address this important question have so far been based on the elimination of the TT network (detubulation) using a hyperosmotic shock with molar concentrations of formamide (13, 14, 16, 17, 20, 21, 27, 41, 50, 70). This technique, first used in skeletal muscle (5), was introduced in the cardiac field by Kawai *et al.* (41) and provided an excellent experimental tool for numerous T-tubular studies. However, the potential role of shock-induced detubulation in physiologically and pathophysiologically relevant conditions is essentially unknown. The method is very harsh on the cells as less than 10% of the cells survive to the procedure, which raises issues about whether the “survivors” are representative cells (12). Besides, osmotic shock produces a sealing of the TT which remain present as vesicles inside the cell (50) and may still respond to hormonal or pharmacological challenges even if they are electrically disconnected from the surface membrane (49).

To overcome these limitations, we introduce here a completely different approach based on size exclusion. We proposed to enlarge the size of a ligand molecule by attaching it to a chain of PolyEthylene Glycol (PEG) so that it does not access TTM but remains active on OSM. PEG is a hydrophilic, flexible and rather inert polymer. The covalent linking of one or several PEG chains to a therapeutic molecule, called PEGylation (73), is commonly used in several products on the market (58). PEGylated molecules exhibit improved biodistribution and pharmacokinetics, better stability and solubility, reduced immunogenicity and longer plasma half-life due to both reduced renal filtration and proteolysis, when compared to non-PEGylated analogues (2, 33). In our study, we diverted the same technology towards an entirely different objective: Instead of using PEGylated molecules to improve drug formulation, pharmacokinetics and efficacy, we propose to use PEGylated drugs as *key holders* to prevent drug (*key*) access in the TT network and thus limit its access to the outer surface of the cell. We provide a proof of concept that this approach works using PEG_5000_ functionalized with non-permeant FITC (PEG-FITC): When ARVMs were exposed to PEG-FITC, fluorescence was only seen on the periphery of the cell while when the non-PEGylated free diffusible dye was used, fluorescence was seen in the TT compartment (Fig. 1). Thus, large molecular weight PEGs are prevented to diffuse within TTs whereas small molecular weight fluorophore can easily enter.

This finding was surprising, though, since longitudinal TT diameter from healthy control ARVMs varies from 50 to 350 nm (~200 nm on average (74)) (67, 74), which is quite larger than the size of a PEG_5000_ molecule on the order of 5 nm. However, a number of recent studies have shown that solute movement in the TT network is strongly restricted even though the diameter of TTs is far larger than the molecular dimensions of typical solutes (26, 37, 43, 60, 72). Impediment of solute movement in the TT network was proposed to be caused by the presence of the glycocalyx (44) at the cell surface, reducing the effective diameter of a TT (37, 43, 56). This was supported by our finding that pre-treatment of isolated ARVMs with neuraminidase, an enzyme which cleaves sialic acid from oligosaccharide chains in the glycoprotein matrix on the external cell surface, allowed PEG-FITC to enter the TT compartment (Fig. 1). Thus, when the protocol used for cell isolation is sufficiently gentle to preserve the integrity of the extracellular matrix in the TT, which must be the case in our experimental conditions, it is possible to prevent a ligand from accessing the TTM if the size of the ligand molecule is sufficiently enlarged. We selected PEG molecules between 3000 and 5000 Da MW which represents a good compromise between solubility issues and size exclusion.

PEG-Iso was thus synthetized by linking isoprenaline to PEG_5000_ molecules. Characterization of its binding properties showed that PEG-Iso binds to β_1_- and β_2_-ARs but with ~2-orders of magnitude lower affinity than Iso (Fig. 2). Since PEG was grafted on the lateral amino group of Iso (Fig. S1), a position shown to maintain its affinity for β-ARs (59), it was unlikely that the chemical modification *per se* was the reason for the decreased affinity. We thus explored a possible involvement of the PEG conformation in this phenomenon using molecular dynamics simulations. Because of its hydrophilicity and flexibility, PEG is very dynamic in solution and constantly changes conformation. Consequently, the Iso moiety in PEG-Iso is rarely fully accessible to solvent and spends most of the time with the PEG chain wrapped around it (Fig. 2). Based on the calculations, only ~1% of PEG-Iso conformations are able to bind β-ARs as compared to 100% for free Iso, which is in the order of magnitude of the measured difference in their respective binding affinity on β-ARs.

When applied on ARVMs, PEG-Iso produced stimulatory effects on [cAMP]_i_, PKA activity, *I*_Ca,L_, sarcomere shortening and Ca^2+^ transients which all had the hallmarks of a β-AR stimulation (Fig. 3 to 8). Because of the reduced binding affinity towards β-ARs, PEG-Iso produced its effect at ~2-orders of magnitude larger concentrations than Iso. But there were two other striking differences between the effects of PEG-Iso and Iso which strongly suggest that the two ligands did not act on the exact same populations of receptors.

First, PEG-Iso produced a stimulation of [cAMP]_i_ with a much lower efficacy than Iso (Fig. 3). The simplest explanation for this result, which is what drove us to undertake this study, is that PEG-Iso activates only β-ARs located in the OSM while Iso activates β-ARs located in both OSM and TTM. Since both populations of β-ARs are likely coupled to ACs (29, 71), one would expect an activation of OSM β-ARs with PEG-Iso to lead to a smaller maximal increase in [cAMP]_i_ than an activation of all β-ARs with Iso. This diminution is in line with decreased cAMP production sites by detubulation (12). Cyclic AMP must emanate mainly from sarcolemmal β_1_-ARs, which produce highly diffusible signals throughout the cell in contrast to β_2_-ARs (52). However, there are conflicting results regarding the distribution of β-ARs on the cardiac cell membrane. Immunohistochemical data have suggested that β_1_- and β_2_-ARs are present in OSM and TTM in mouse heart (78). Using radioligand binding, β_1_-AR density was found to be almost 2-fold more concentrated in OSM that TTM, whereas β_2_-AR density was evenly distributed across the entire cell surface in dog heart (35). Using nanoscale live-cell scanning ion conductance and fluorescence resonance energy transfer microscopy techniques in healthy ARVMs, Nikolaev *et al.* found that β_1_-ARs are distributed across the entire cell surface while β_2_-ARs are localized exclusively in TTM (53). While the distribution of β_1_-ARs across the entire cell surface was confirmed in ARVMs in another study using formamide detubulation, β_2_-ARs were found to be only present in OSM (20). Additional experiments using PEGylated selective β_1_- and β_2_-AR agonists and antagonists are needed to solve this issue and to precisely characterize the role of each β-AR subtype in the different membrane compartments.

Second, PEG-Iso produced a much lower stimulation of nuclear PKA activity than Iso (Fig. 8) even though both ligands were equally efficient in stimulating cytosolic PKA activity (Fig. 7). Also, when cytosolic and nuclear PKA activity measured during PEG-Iso or Iso stimulation were plotted as a function of [cAMP]_i_ measured under the same conditions, PEG-Iso was found to be more efficient than Iso to increase PKA activity in the cytosol, while on the contrary Iso is more efficient than PEG-Iso to increase PKA activity in the nucleus (Fig. S3). A likely interpretation of these results is that β-ARs located in TTM (or a β-AR subtype more concentrated in TTM) are more efficiently coupled to nuclear PKA than those located in OSM, while β-ARs located in OSM (or a β-AR subtype more concentrated in OSM) are more efficiently coupled to cytosolic PKA. In a recent study, we showed that β_2_-ARs, unlike β_1_-ARs, are inefficient in activating nuclear PKA activity in ARVMs (6). Thus, if the ratio of β_2_/β_1_ receptors is larger in OSM than in TTM, as discussed above (20), then this would explain the lower efficacy of PEG-Iso to activate nuclear PKA activity as compared to Iso. Recent studies have shown that perinuclear PKA has the ability to activate without detachment of the catalytic subunits, and thus to phosphorylate its targets in close proximity (65, 66). According to these studies, only supra-physiological concentrations of cAMP could lead to dissociation of the catalytic subunits and their translocation into the nucleus. Perinuclear PKA is anchored to the nuclear membrane via its interaction with mAKAP and its dynamics are finely regulated by AC5 and PDE4D3 (22), AC5 being concentrated in TTM (71) and PDE4D3 being linked to mAKAP (23). The location of AC5 may therefore be crucial for nuclear signalling and explain why β-ARs present in TTM are necessary for activation of nuclear PKA.

Another important finding from this study concerns the contribution of OSM β-ARs to the regulation of EC coupling. Activation of β-ARs only in OSM with PEG-Iso produced a similar increase in *I*_Ca,L_ amplitude, sarcomere shortening and Ca^2+^ transients as Iso, when both ligands were used at concentrations producing an equivalent elevation of [cAMP]_i_. Since OSM contributes to ~60% of total cell membrane in ARVMs (12), either β-ARs and ACs are more concentrated in OSM than TTM, or they are in large excess over what is needed to activate PKA phosphorylation of proteins involved in EC coupling. Also, cAMP produced at OSM must diffuse rapidly in the cytosol in order to activate PKA phosphorylation of substrates located deep inside the cell, such as LTCCs in TTM. A recent study showed that under basal conditions, cAMP is bound to proteins and its diffusion in the cytosol is slow; but when cAMP increases in the cytosol, e.g. as a result of β-AR stimulation, the cAMP binding sites become saturated and cAMP diffuses freely (10).

While this study provides new insight on the differential function of OSM vs. TTM β-ARs in healthy cardiomyocytes, it raises obvious questions on how their role is altered in cardiac pathology. A reduction in the density and a disorganization of the TT network was observed in pathological conditions (31, 74), with great impact on EC coupling and contractile function (63). A previous study showed a redistribution of β_2_-ARs from TTM to OSM in failing ARVMs (53). As β-ARs are not simply bystanders but also participate in the remodeling during pathological hypertrophy and failure, it is important to know how the OSM *vs.* TTM subpopulations of β-AR subtypes contribute to this process. The PEGylation technology should allow to elucidate the changes that operate in OSM *vs.* TTM distribution of β-ARs in ventricular cells isolated from diseased rats, and to determine how this impacts on cellular function.

In conclusion, this study provides a novel approach to distinguish the function of a membrane protein depending on its location on the cell membrane. Whereas we focused here on the function of β-ARs, the size exclusion strategy provided by ligand PEGylation can be extended to other ligands, such as selective agonists or antagonists of β-AR subtypes or other G-protein coupled receptors, dihydropyridine agonists or antagonists of LTCCs, etc. More generally, it should pave the road to exploring and comparing the function of any membrane receptor, channel, transporter or enzyme in TTM and OSM compartments in intact cardiomyocytes.

## Materials and Methods

### Animals

All animal care and experimental procedures complied with the ARRIVE guidelines and conform to the European Community guiding principles in the Care and Use of Animals (Directive 2010/63/EU of the European Parliament), the local Ethics Committee (CREEA Ile-de France Sud) guidelines, and the French decree no. 2013-118 of February 1st, 2013 on the protection of animals used for scientific purposes (JORF no. 0032, February 7th, 2013, p2199, text no. 24). Animal experiments according were carried out according to the European Community guiding principles in the care and use of animals (2010/63/UE), the local Ethics Committee (CREEA Ile-de-France Sud) guidelines and the French decree n° 2013-118 on the protection of animals used for scientific purposes.

### Synthesis of PEG-ISO

Isoprenaline was grafted into the reactive end of 5000 Da PEG (PEG_5000_). A carbodiimide reaction was carried out between the carboxylic acid of PEG and the amine function of Iso (Supplementay Fig. 1). In a 25 mL amber vial, 41.3 mg of DCC (0.2 mmol; 2eq), 23 mg of NHS (0.2 mmol; 2eq) and 500 mg of OH-PEG_5000_-COOH (0.1 mmol; 1eq) were solubilized in 10 mL of dichloromethane. 1 mL of triethylamine (TEA) was added to the syringe. The solution was stirred for 5 h at RT. 21.1 mg isoprenaline (0.1 mmol; 1 eq) were added to the reaction mixture. The mixture was left to stir overnight. The solution was then filtered through silica gel and precipitated twice in cold diethyl ether. After evaporation of the ether, the final powder was solubilised in H_2_O.

### Purification of PEG-ISO

A purification step was introduced to obtain a pure PEG-Iso product and to eliminate the various by-products obtained, potential impurities and especially traces of ungrafted Iso which could interfere in the biological experiments. An HPLC 1290 Infinity II - (Agilent Technologies) consisting of a 4-channel binary pump, a UV/visible detector (PDA diode array), a 1260 Infinity DEDL light scattering detector and a C18 XBridge column, 4.6 x 150 mm, 5 µm (17 mL/min flow rate) was used. The mobile phase consisted of a mixture of water + 0.1% formic acid and acetonitrile. This phase was pushed according to a gradient from 1 to 100% in 15 min into the stationary phase for the purification of PEG-Iso. Detection was carried out in the UV at 254 and 280 nm. PEG-Iso was solubilized in water. Several successive injections of 300 µL each were performed. The retention time of PEG-Iso was 11 min. The fractions containing PEG-Iso were recovered, evaporated and then frozen and freeze-dried. Finally, the powder obtained was stored at −20°C and protected from light.

### Cardiomyocyte isolation and culture

Adult male Wistar rats (250–300 g) were anesthetized by intraperitoneal injection of pentobarbital (0.1 mg/g) and hearts were excised rapidly. Individual adult rat ventricular cardiomyocytes were obtained by retrograde perfusion of the heart as previously described (48). For enzymatic dissociation, the hearts were perfused during 5 min at a constant flow of 6 mL/min at 37°C with a Ca^2+^-free Ringer solution containing (in mM): NaCl 117, KCl 5.7, NaHCO_3_ 4.4, KH_2_PO_4_ 1.5, MgCl_2_ 1.7, D-glucose 11.7, Na_2_-phosphocreatine 10, taurine 20, and 4-(2-hydroxyethyl)piperazine-1-ethanesulfonic acid (HEPES) 21, pH 7.1. This was followed by a perfusion at 4 mL/min for 40 min with the same solution containing 1 mg/mL of collagenase A (Roche Diagnostics GmbH, Mannheim, Germany) plus 300 µM ethylene glycol tetraacetic acid (EGTA) and CaCl_2_ to adjust free Ca^2+^ concentration to 20 µM. The ventricles were then separated, finely chopped and gently agitated to dissociate individual cells. The resulting cell suspension was filtered on gauze and the cells were allowed to settle down. The supernatant was discarded and cells resuspended four more times with Ringer solution at increasing [Ca^2+^] from 20 to 300 µM. Freshly isolated cells were suspended in minimal essential medium (MEM: M 4780; Sigma, St Louis, MO USA) containing 1.2 mM [Ca^2+^] supplemented with 2.5% foetal bovine serum (FBS, Invitrogen, Cergy-Pontoise, France), 1% penicillin–streptomycin (P-S), 20 mM HEPES (pH 7.6), and plated on 35 mm, laminin-coated (10 mg/mL) culture dishes at a density of 10^4^ cells per dish and kept at 37°C (5% CO_2_). After 1 h, the medium was replaced by 300 µL of FBS-free MEM. To perform FRET imaging, the medium was replaced by 300 μL of FCS-free MEM or transduced with an adenovirus encoding the Epac-S^H187^ FRET-based sensor(42), or the AKAR3 FRET-based sensor(3) addressed to the cytosol or the nucleus, at a multiplicity of infection (MOI) of 1000 pfu/cell and cells were cultivated for 36 h prior to the experiments. Patch-clamp and IonOptix experiments were performed on cells 24 h after dissociation.

### Confocal imaging

Ventricular cardiomyocytes from adult rats were deposited on 35 mm glass bottom Petri dishes. Two hours after deposition, the control cells were brought into contact with commercial fluorescent PEG_5000_ functionalized with FITC (Fluorescein Isothiocyanate) (Nanocs) or free Fluorescein (Sigma-Aldrich) chosen as positive control. Other cells were treated with neuraminidase during 1 h at 0.25 U/mL. After that, cells were incubated with PEG-FITC. All molecules were dissolved in Ringer containing (in mM): NaCl 121.6, KCl 5.4, MgCl_2_ 1.8, CaCl_2_ 1.8, NaHCO_3_ 4, NaH_2_PO_4_ 0.8, D-glucose 5, sodium pyruvate 5, HEPES 10, adjusted to pH 7.4. The acquisitions were made with a Leica TCS SP5 confocal microscope, a white light laser and an X40 oil immersion objective. The cells were excited at 495 nm and fluorescence was recovered at wavelengths >510 nm.

### Molecular dynamics simulations

Molecular dynamics simulations of free Iso and PEG-Iso were carried out to predict their conformations in aqueous solution and in particular to assess the solvent-accessibility of Iso moiety once grafted to the PEG. A 3000 Da PEG molecule was used in these simulations. The 2D chemical structure of each of the two molecules was first converted into a three-dimensional structure using ChemAxon’s MarvinSketch chemical editing software. The topology and GAFF force field parameters (75) used in this study were then automatically generated using the ACPYPE program (69). Initial three-dimensional structure was then placed in the centre of 13.4 nm side cubic simulation. These dimensions allow the solute not to interact with its virtually created images due to the periodic boundary conditions used to simulate an infinitely duplicated mesh. The box was then filled with water (model TIP3P(39)) and 150 mM NaCl. The system was then subjected to 2 short simulations (1 ns each) to equilibrate first the temperature around 300 K and then the pressure around 1 bar. The Nosé-Hoover (38, 54) (coupling time τT = 0.5 ps) and Parrinello-Rahman (57) (τP = 2.5 ps) algorithms were used for maintaining constant temperature and pressure, respectively. Finally, each of the two molecules was subjected to a production run of 150 ns. All simulations were performed using the GROMACS software version 2016(1). To analyse the trajectories, we mainly used the GROMACS internal *gmx sasa* tool to calculate the solvent-accessible surface area (SASA) for Iso moiety in both free Iso or PEG-Iso molecules. To compare the solvent-accessibility of Iso moiety in free Iso and PEG-Iso, the percentage of PEG-Iso conformations with a SASA_PEG-Iso_ ≥ 0.9 SASA_free-Iso_ was calculated.

### Binding experiments

Binding assays were carried out in a final volume of 100 μL, containing membrane suspension fro CHO cells overexpressing human ß_1_- or ß_2_-adrenergic receptors, 145 pM of the radioligand [^125^Iodo]cyanopindolol and non-radioactive ligands at concentrations ranging from 10^−10^ to 10^−3^ M. Incubation was carried out for 2 h at room temperature and terminated by addition of 4 mL PBS. Then, rapid filtration was performed through Whatman GF/C glass fiber filters previously soaked in PBS containing 0.33% polyethyleneimine using a compact cell harvester (Millipore^®^ 1225 Sampling Vacuum Manifold). Binding reaction was transferred to the filters, and washed 3 times with Wash Buffer. Filter-bound radioactivity was measured in a gamma counter. All determinations were performed at least in triplicates.

### FRET imaging

FRET experiments were performed at room temperature 36 h after cell plating. Cells were maintained in a Ringer solution containing (in mM): NaCl 121.6, KCl 5.4, MgCl_2_ 1.8; CaCl_2_ 1.8; NaHCO_3_ 4, NaH_2_PO_4_ 0.8, D-glucose 5, sodium pyruvate 5, HEPES 10, adjusted to pH 7.4. Images were captured every 4 s using the 40x oil immersion objective of a Nikon TE 300 inverted microscope connected to a software-controlled (Metafluor, Molecular Devices, Sunnyvale, CA, USA) cooled charge coupled (CCD) camera (Sensicam PE, PCO, Kelheim, Germany). Cells were excited during 150-300 ms by a Xenon lamp (100 W, Nikon, Champigny-sur-Marne, France) using a 440/20BP filter and a 455LP dichroic mirror. Dual emission imaging was performed using an Optosplit II emission splitter (Cairn Research, Faversham, UK) equipped with a 495LP dichroic mirror and BP filters 470/30 (CFP) and 535/30 (YFP), respectively. Spectral bleed-through into the YFP channel was subtracted using the formula: YFP_corr_=YFP-0.6xCFP.

### Electrophysiological experiments

The whole cell configuration of the patch-clamp technique was used to record *I*_Ca,L_. Patch electrodes had resistance of 0.5–1.5 MΩ when filled with internal solution containing (in mM): CsCl 118, EGTA 5, MgCl_2_ 4, Na_2_-phosphocreatine 5, Na_2_ATP 3.1, Na_2_GTP 0.42, CaCl_2_ 0.062, Hepes 10, adjusted to pH 7.3 with CsOH. External Cs^+^-Ringer solution contained (in mM): NaCl 107.1, CsCl 20, NaHCO_3_ 4, NaH_2_PO_4_ 0.8, D-glucose 5, sodium pyruvate 5, Hepes 10, MgCl_2_ 1.8, CaCl_2_ 1.8, adjusted to pH 7.4 with NaOH. The cells were depolarized every 8 s from –50 mV to 0 mV during 400 ms. The junction potential was adjusted to give zero current between pipette and bath solution before the cells were attached to obtain a tight gigaseal (>1GΩ). The use of −50mV as holding potential allowed the inactivation of voltage-dependent sodium currents. Currents were not compensated for capacitance and leak currents. The amplitude of *I*_Ca,L_ was measured as the difference between the peak inward current and the current at the end of the depolarization pulses. The cells were voltage-clamped a patch-clamp amplifier (either RK-400 (Bio-Logic,Claix, France) or Axopatch 200B (Axon Instruments, Inc., UnionCity, CA, USA)). Currents were analog filtered at 5 kHz and digitally sampled at 10 kHz using a 16-bit analog to digital converter (DT321; Data translation, Marlboro, MA, USA or a Digidata1440A, Axon Instruments) connected to a PC (Dell, Austin, TX, USA).

### Measurements of Ca^2+^ transients and sarcomere shortening

All experiments were performed at room temperature. Isolated ARVMs were loaded with 1 µM Fura-2 AM (Thermo Fischer Scientific) and Pluronic acid (0.012%, Thermo Fischer Scientific) for 15 min in a Ringer solution containing (in mM): KCl 5.4, NaCl 121.6, sodium pyruvate 5, NaHCO_3_ 4.013, NaH_2_PO_4_ 0.8, CaCl_2_ 1.8, MgCl_2_ 1.8, glucose 5, Hepes 10, pH 7.4 with NaOH. Sarcomere shortening and Fura-2 ratio (measured at 512 nm upon excitation at 340 and 380 nm) were simultaneously recorded in Ringer solution, using a double excitation spectro fluorimeter coupled with a video detection system (IonOptix, Milton, MA, USA). Myocytes were electrically stimulated, with biphasic field pulses (5 V, 4 ms) at a frequency of 1 Hz. Ca^2+^ transient amplitude was measured by dividing the twitch amplitude (difference between the end-diastolic and the peak systolic ratios) by the end-diastolic ratio, thus corresponding to the percentage of variation in the Fura-2 ratio. Similarly, sarcomere shortening was assessed by its percentage variation, which was obtained by dividing the twitch amplitude (difference between the end-diastolic and the peak systolic sarcomere length) by the end-diastolic sarcomere length. Relaxation kinetics were estimated by a non-linear fit of the decaying part of the Ca^2+^ transient and sarcomere shortening traces with the following equation: X(t) = A.exp(−(t−t_0_)/τ)+A_0_, where t_0_ is zero time, A_0_ the asymptote of the exponential, A the relative amplitude of the exponential and τ the time constant of relaxation. The maximum first derivative of transients during the deflexion allowed determination of the rising velocities of the signals. All parameters were calculated offline with dedicated software (IonWizard 6x, IonOptix R).

### Proteins expression analysis

Cardiomyocytes were treated with or without isoprenaline 10 nM or PEG-Iso 1 µM (for 15 min) and the nuclei were enriched using a purification kit (proteoExtract). Nuclear proteins were examined by immunoblot for phosphorylation of PKA substrates (with a specific phospho-PKA antibody) and total protein. The ratio of phospho-PKA expression was assessed by densimetric scanning of immunoblots.

### Statistical analysis

All results are expressed as mean±SEM. Statistical analysis was performed using GraphPad software (GraphPad Software, Inc., La Jolla, CA, USA). Normal distribution was tested by the Shapiro-Wilk normality test. For simple two-group comparison, we used an unpaired Student *t* test. Differences between multiple groups were analysed using an ordinary 1-way ANOVA with Tukey post hoc test. A P value<0.05 was considered statistically significant.

## Acknowledgements

MB was supported by doctoral grant from the Laboratory of Excellence LERMIT supported by the French National Research Agency (ANR-10-LABX-33) under the program “Investissements d’Avenir” ANR-11-IDEX-0003-01. She also received a doctoral grant from the Fondation pour la Recherche Médicale. This work was also funded by grant ANR-15-CE14-0014-01 to RF. Authors would like to thank K. Leblanc (BioCIS) for help with preparative HPLC.

## References

1. Abraham MJ, Murtola T, Schulz R, Páll S, Smith JC, Hess B, Lindahl E. GROMACS: High performance molecular simulations through multi-level parallelism from laptops to supercomputers. SoftwareX 1–2: 19–25, 2015.

2. Abuchowski A, van Es T, Palczuk NC, Davis FF. Alteration of immunological properties of bovine serum albumin by covalent attachment of polyethylene glycol. J Biol Chem 252: 3578–3581, 1977.

3. Allen MD and Zhang J. Subcellular dynamics of protein kinase A activity visualized by FRET-based reporters. Biochemical and biophysical research communications 348: 716–721, 2006.

4. Antos CL, Frey N, Marx SO, Reiken S, Gaburjakova M, Richardson JA, Marks AR, Olson EN. Dilated cardiomyopathy and sudden death resulting from constitutive activation of protein kinase A. Circulation research 89: 997–1004, 2001.

5. Argiro V. Excitation-contraction uncoupling of striated muscle fibres by formamide treatment: evidence of detubulation. Journal of muscle research and cell motility 2: 283–294, 1981.

6. Bedioune I, Lefebvre F, Lechêne P, Varin A, Domergue V, Kapiloff MS, Fischmeister R, Vandecasteele G. PDE4 and mAKAPβ are nodal organizers of β_2_-AR nuclear PKA signaling in cardiac myocytes. Cardiovascular research 114: 1499–1511, 2018.

7. Berry RG, Despa S, Fuller W, Bers DM, Shattock MJ. Differential distribution and regulation of mouse cardiac Na^+^/K^+^-ATPase α_1_ and α_2_ subunits in T-tubule and surface sarcolemmal membranes. Cardiovascular research 73: 92–100, 2007.

8. Bers DM. Calcium cycling and signaling in cardiac myocytes. Ann Rev Physiol 70: 23–49, 2008.

9. Bers DM. Cardiac excitation-contraction coupling. Nature 415: 198–205, 2002.

10. Bock A, Annibale P, Konrad C, Hannawacker A, Anton SE, Maiellaro I, Zabel U, Sivaramakrishnan S, Falcke M, Lohse MJ. Optical mapping of cAMP signaling at the nanometer scale. Cell 182: 1519–1530, 2020.

11. Boluyt MO, Oneill L, Meredith AL, Bing OHL, Brooks WW, Conrad CH, Crow MT, Lakatta EG. Alterations in cardiac gene expression during the transition from stable hypertrophy to heart failure -Marked upregulation of genes encoding extracellular matrix components. Circulation research 75: 23–32, 1994.

12. Bourcier A, Barthe M, Bedioune I, Lechene P, Miled HB, Vandecasteele G, Fischmeister R, Leroy J. Imipramine as an alternative to formamide to detubulate rat ventricular cardiomyocytes. Experimental physiology 104: 1237–1249, 2019.

13. Brette F, Despa S, Bers DM, Orchard CH. Spatiotemporal characteristics of SR Ca^2+^ uptake and release in detubulated rat ventricular myocytes. J Mol Cell Cardiol 39: 804–812, 2005.

14. Brette F, Komukai K and Orchard CH. Validation of formamide as a detubulation agent in isolated rat cardiac cells. American journal of physiology Heart and circulatory physiology 283: H1720–H1728, 2002.

15. Brette F and Orchard C. T-tubule function in mammalian cardiac myocytes. Circulation research 92: 1182–1192, 2003.

16. Brette F, Salle L and Orchard CH. Differential modulation of L-type Ca^2+^ current by SR Ca^2+^ release at the T-tubules and surface membrane of rat ventricular myocytes. Circulation research 95: e1–7, 2004.

17. Brette F, Salle L and Orchard CH. Quantification of calcium entry at the T-tubules and surface membrane in rat ventricular myocytes. Biophys J 90: 381–389, 2006.

18. Caldwell JL, Smith CE, Taylor RF, Kitmitto A, Eisner DA, Dibb KM, Trafford AW. Dependence of cardiac transverse tubules on the BAR domain protein amphiphysin II (BIN-1). Circulation research 115: 986–996, 2014.

19. Carl SL, Felix K, Caswell AH, Brandt NR, Ball WJ, Vaghy PL, Meissner G, Ferguson DG. Immunolocalization of sarcolemmal dihydropyridine receptor and sarcoplasmic reticular triadin and ryanodine receptor in rabbit ventricle and atrium. The Journal of cell biology 129: 673–682, 1995.

20. Cros C and Brette F. Functional subcellular distribution of β_1_- and β_2_-adrenergic receptors in rat ventricular cardiac myocytes. Physiol Rep 1: e00038, 2013.

21. Despa S, Brette F, Orchard CH, Bers DM. Na/Ca exchange and Na/K-ATPase function are equally concentrated in transverse tubules of rat ventricular myocytes. Biophys J 85: 3388–3396, 2003.

22. Diviani D, L Dodge-Kafka K, Li J, Kapiloff MS. A-kinase anchoring proteins: Scaffolding proteins in the heart. American journal of physiology Heart and circulatory physiology 301: H1742–H1753, 2011.

23. Dodge-Kafka KL, Soughayer J, Pare GC, Carlisle Michel JJ, Langeberg LK, Kapiloff MS, Scott JD. The protein kinase A anchoring protein mAKAP co-ordinates two integrated cAMP effector pathways. Nature 437: 574–578, 2005.

24. Eisner DA, Caldwell JL, Kistamas K, Trafford AW. Calcium and excitation-contraction coupling in the heart. Circulation research 121: 181–195, 2017.

25. Engelhardt S, Hein L, Wiesmann F, Lohse MJ. Progressive hypertrophy and heart failure in β_1_-adrenergic receptor transgenic mice. Proc Natl Acad Sci USA 96: 7059–7064, 1999.

26. Entcheva E. Uncovering an electrically heterogeneous cardiomyocyte by FRAP-quantified diffusion in the T-tubules. Proc Natl Acad Sci USA 115: E560–E561, 2018.

27. Fowler MR, Dobson RS, Orchard CH, Harrison SM. Functional consequences of detubulation of isolated rat ventricular myocytes. Cardiovascular research 62: 529–537, 2004.

28. Fu Y, Shaw S, Naami R, Vuong C, Basheer WA, Guo X, Hong T. Isoproterenol promotes rapid ryanodine receptor movement to BIN1 organized dyads. Circulation 133: 388–397, 2016.

29. Gao TY, Puri TS, Gerhardstein BL, Chien AJ, Green RD, Hosey MM. Identification and subcellular localization of the subunits of L-type calcium channels and adenylyl cyclase in cardiac myocytes. J Biol Chem 272: 19401–19407, 1997.

30. Gorelik J, Wright PT, Lyon AR, Harding SE. Spatial control of the βAR system in heart failure: the transverse tubule and beyond. Cardiovascular research 98: 216–224, 2013.

31. Guo A, Zhang C, Wei S, Chen B, Song LS. Emerging mechanisms of T-tubule remodelling in heart failure. Cardiovascular research 98: 204–215, 2013.

32. Haj Slimane Z, Bedioune I, Lechêne P, Varin A, Lefebvre F, Mateo P, Domergue-Dupont V, Dewenter M, Richter W, Conti M, El-Armouche A, Zhang J, Fischmeister R, Vandecasteele G. Control of cytoplasmic and nuclear protein kinase A activity by phosphodiesterases and phosphatases in cardiac myocytes. Cardiovascular research 102: 97–106, 2014.

33. Harris JM and Chess RB. Effect of pegylation on pharmaceuticals. Nat Rev Drug Discov 2: 214–221, 2003.

34. Hatano A, Okada J, Hisada T, Sugiura S. Critical role of cardiac t-tubule system for the maintenance of contractile function revealed by a 3D integrated model of cardiomyocytes. Journal of biomechanics 45: 815–823, 2012.

35. He JQ, Balijepalli RC, Haworth RA, Kamp TJ. Crosstalk of β-adrenergic receptor subtypes through G_i_ blunts beta-adrenergic stimulation of L-type Ca^2+^ channels in canine heart failure. Circulation research 97: 566–573, 2005.

36. Hong T and Shaw RM. Cardiac T-tubule microanatomy and function. Physiological reviews 97: 227–252, 2017.

37. Hong T, Yang H, Zhang SS, Cho HC, Kalashnikova M, Sun B, Zhang H, Bhargava A, Grabe M, Olgin J, Gorelik J, Marban E, Jan LY, Shaw RM. Cardiac BIN1 folds T-tubule membrane, controlling ion flux and limiting arrhythmia. Nat Med 20: 624–632, 2014.

38. Hoover WG. Canonical dynamics: Equilibrium phase-space distributions. Phys Rev A Gen Phys 31: 1695–1697, 1985.

39. Jorgensen WL, Chandrasekhar J, Madura JD, Impey RW, Klein ML. Comparison of simple potential functions for simulating liquid water. J Chemical Physics 79: 926–935, 1983.

40. Karam S, Margaria JP, Bourcier A, Mika D, Varin A, Bedioune I, Lindner M, Bouadjel K, Dessillons M, Gaudin F, Lefebvre F, Mateo P, Lechene P, Gomez S, Domergue V, Robert P, Coquard C, Algalarrondo V, Samuel JL, Michel JB, Charpentier F, Ghigo A, Hirsch E, Fischmeister R, Leroy J, Vandecasteele G. Cardiac overexpression of PDE4B blunts β-adrenergic response and maladaptive remodeling in heart failure. Circulation 142: 161–174, 2020.

41. Kawai M, Hussain M and Orchard CH. Excitation-contraction coupling in rat ventricular myocytes after formamide-induced detubulation. Am J Physiol 277: H603–609, 1999.

42. Klarenbeek J, Goedhart J, van Batenburg A, Groenewald D, Jalink K. Fourth-generation EPAC-based fret sensors for cAMP feature exceptional brightness, photostability and dynamic range: Characterization of dedicated sensors for flim, for ratiometry and with high affinity. PloS one 10: e0122513, 2015.

43. Kong CHT, Rog-Zielinska EA, Kohl P, Orchard CH, Cannell MB. Solute movement in the t-tubule system of rabbit and mouse cardiomyocytes. Proc Natl Acad Sci USA 115: E7073–E7080, 2018.

44. Langer GA, Frank JS and Philipson KD. Ultrastructure and calcium exchange of the sarcolemma, sarcoplasmic reticulum and mitochondria of the myocardium. Pharmacology & therapeutics 16: 331–376, 1982.

45. Leroy J and Fischmeister R. β-adrenergic regulation of the L-type Ca^2+^ current: The missing link eventually discovered. Med Sci (Paris) 36: 569–572, 2020.

46. Leroy J, Vandecasteele G and Fischmeister R. Cyclic AMP signaling in cardiac myocytes. Curr Op Physiol 1: 161–171, 2018.

47. Liu G, Papa A, Katchman AN, Zakharov SI, Roybal D, Hennessey JA, Kushner J, Yang L, Chen BX, Kushnir A, Dangas K, Gygi SP, Pitt GS, Colecraft HM, Ben-Johny M, Kalocsay M, Marx SO. Mechanism of adrenergic CaV1.2 stimulation revealed by proximity proteomics. Nature 577: 695–700, 2020.

48. Mika D, Bobin P, Pomérance M, Lechêne P, Westenbroek R, Catterall WA, Vandecasteele G, Leroy J, Fischmeister R. Differential regulation of cardiac excitation-contraction coupling by cAMP phosphodiesterase subtypes. Cardiovascular research 100: 336–346, 2013.

49. Moench I and Lopatin AN. Ca^2+^ homeostasis in sealed t-tubules of mouse ventricular myocytes. J Mol Cell Cardiol 72: 374–383, 2014.

50. Moench I, Meekhof KE, Cheng LF, Lopatin AN. Resolution of hyposmotic stress in isolated mouse ventricular myocytes causes sealing of t-tubules. Experimental physiology 98: 1164–1177, 2013.

51. Morisco C, Zebrowski DC, Vatner DE, Vatner SF, Sadoshima J. ß-Adrenergic cardiac hypertrophy is mediated primarily by the ß_1_-subtype in the rat heart. J Mol Cell Cardiol 33: 561–573, 2001.

52. Nikolaev VO, Bunemann M, Schmitteckert E, Lohse MJ, Engelhardt S. Cyclic AMP imaging in adult cardiac myocytes reveals far-reaching ß_1_-adrenergic but locally confined ß_2_-adrenergic receptor-mediated signaling. Circulation research 99: 1084–1091, 2006.

53. Nikolaev VO, Moshkov A, Lyon AR, Miragoli M, Novak P, Paur H, Lohse MJ, Korchev YE, Harding SE, Gorelik J. ß_2_-Adrenergic receptor redistribution in heart failure changes cAMP compartmentation. Science 327: 1653–1657, 2010.

54. Nosé S. A unified formulation of the constant temperature molecular dynamics methods. J Chemical Physics 81: 511–519, 1984.

55. Osadchii OE. Myocardial phosphodiesterases and regulation of cardiac contractility in health and cardiac disease. Cardiovasc Drugs Ther 21: 171–194, 2007.

56. Parfenov AS, Salnikov V, Lederer WJ, Lukyanenko V. Aqueous diffusion pathways as a part of the ventricular cell ultrastructure. Biophys J 90: 1107–1119, 2006.

57. Parrinello M and Rahman A. Polymorphic transitions in single crystals: A new molecular dynamics method. J Applied Physics 52: 7182–7190 1981.

58. Pasut G and Veronese FM. State of the art in PEGylation: the great versatility achieved after forty years of research. J Control Release 161: 461–472, 2012.

59. Ruffolo RR, Bondinell W and Hieble JP. α-and β-Adrenoceptors: From the gene to the clinic. 2. Structure-activity relationships and therapeutic applications. Journal of medicinal chemistry 38: 3681–3716, 1995.

60. Scardigli M, Crocini C, Ferrantini C, Gabbrielli T, Silvestri L, Coppini R, Tesi C, Rog-Zielinska EA, Kohl P, Cerbai E, Poggesi C, Pavone FS, Sacconi L. Quantitative assessment of passive electrical properties of the cardiac T-tubular system by FRAP microscopy. Proc Natl Acad Sci USA 114: 5737–5742, 2017.

61. Schobesberger S, Wright PT, Poulet C, Sanchez Alonso Mardones JL, Mansfield C, Friebe A, Harding SE, Balligand JL, Nikolaev VO, Gorelik J. β_3_-Adrenoceptor redistribution impairs NO/cGMP/PDE2 signalling in failing cardiomyocytes. Elife 9: e52221, 2020.

62. Scriven DRL, Dan P and Moore EDW. Distribution of proteins implicated in excitation-contraction coupling in rat ventricular myocytes. Biophys J 79: 2682–2691, 2000.

63. Sipido KR and Cheng H. T-tubules and ryanodine receptor microdomains: on the road to translation. Cardiovascular research 98: 159–161, 2013.

64. Skeberdis VA, Gendvilienë V, Zablockaitë D, Treinys R, Maèianskienë R, Bogdelis A, Jurevièius J, Fischmeister R. ß_3_-adrenergic receptor activation increases human atrial tissue contractility and stimulates the L-type Ca^2+^ current. The Journal of clinical investigation 118: 3219–3227, 2008.

65. Smith FD, Esseltine JL, Nygren PJ, Veesler D, Byrne DP, Vonderach M, Strashnov I, Eyers CE, Eyers PA, Langeberg LK, Scott JD. Local protein kinase A action proceeds through intact holoenzymes. Science 356: 1288–1293, 2017.

66. Smith FD, Reichow SL, Esseltine JL, Shi D, Langeberg LK, Scott JD, Gonen T. Intrinsic disorder within an AKAP-protein kinase A complex guides local substrate phosphorylation. Elife 2: e01319, 2013.

67. Soeller C and Cannell MB. Examination of the transverse tubular system in living cardiac rat myocytes by 2-photon microscopy and digital image-processing techniques. Circulation research 84: 266–275, 1999.

68. Song LS, Sobie EA, McCulle S, Lederer WJ, Balke CW, Cheng H. Orphaned ryanodine receptors in the failing heart. Proc Natl Acad Sci USA 103: 4305–4310, 2006.

69. Sousa da Silva AW and Vranken WF. ACPYPE - AnteChamber PYthon Parser interfacE. BMC research notes 5: 367, 2012.

70. Thomas MJ, Sjaastad I, Andersen K, Helm PJ, Wasserstrom JA, Sejersted OM, Ottersen OP. Localization and function of the Na^+^/Ca^2+^-exchanger in normal and detubulated rat cardiomyocytes. J Mol Cell Cardiol 35: 1325–1337, 2003.

71. Timofeyev V, Myers RE, Kim HJ, Woltz R, Sirish P, Heiserman J, Li N, Singapuri A, Tang T, Yarov-Yarovoy V, Yamoah EN, Hammond K, Chiamvimonvat N. Adenylyl cyclase subtype-specific compartmentalization: Differential regulation of L-type Ca^2+^ current in ventricular myocytes. Circulation research 112: 1567–1576, 2013.

72. Uchida K and Lopatin AN. Diffusional and electrical properties of T-tubules are governed by their constrictions and dilations. Biophys J 114: 437–449, 2018.

73. Veronese FM and Harris JM. Introduction and overview of peptide and protein pegylation. Adv Drug Deliv Rev 54: 453–456, 2002.

74. Wagner E, Lauterbach MA, Kohl T, Westphal V, Williams GS, Steinbrecher JH, Streich JH, Korff B, Tuan HT, Hagen B, Luther S, Hasenfuss G, Parlitz U, Jafri MS, Hell SW, Lederer WJ, Lehnart SE. Stimulated emission depletion live-cell super-resolution imaging shows proliferative remodeling of T-tubule membrane structures after myocardial infarction. Circulation research 111: 402–414, 2012.

75. Wang J, Wolf RM, Caldwell JW, Kollman PA, Case DA. Development and testing of a general amber force field. J Comput Chem 25: 1157–1174, 2004.

76. Xiang YK. Compartmentalization of β-adrenergic signals in cardiomyocytes. Circulation research 109: 231–244, 2011.

77. Yang Z, Pascarel C, Steele DS, Kornukai K, Brette F, Orchard CH. Na^+^-Ca^2+^ exchange activity is localized in the T-tubules of rat ventricular myocytes. Circulation research 91: 315–322, 2002.

78. Zhou YY, Yang D, Zhu WZ, Zhang SJ, Wang DJ, Rohrer DK, Devic E, Kobilka BK, Lakatta EG, Cheng HP, Xiao RP. Spontaneous activation of β_2_-but not β_1_-adrenoceptors expressed in cardiac myocytes from β_1_β_2_ double knockout mice. Molecular pharmacology 58: 887–894, 2000.

