## Supplemental Figures S1-S2-S3 for "Distinct functions of cardiac β-adrenergic receptors in the T-tubule *vs.* outer surface membrane"

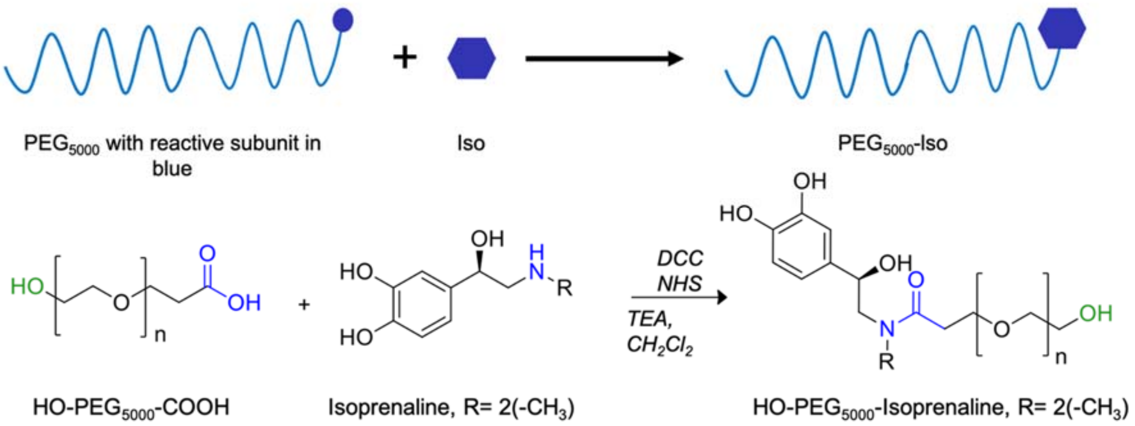


**Fig. S1. PEG-Isoprenaline synthesis. Top**, Concept of chemical grafting between $PEG_{5000}$ and Iso; **Bottom**, chemical coupling between PEG and Iso using carbodiimide reaction.

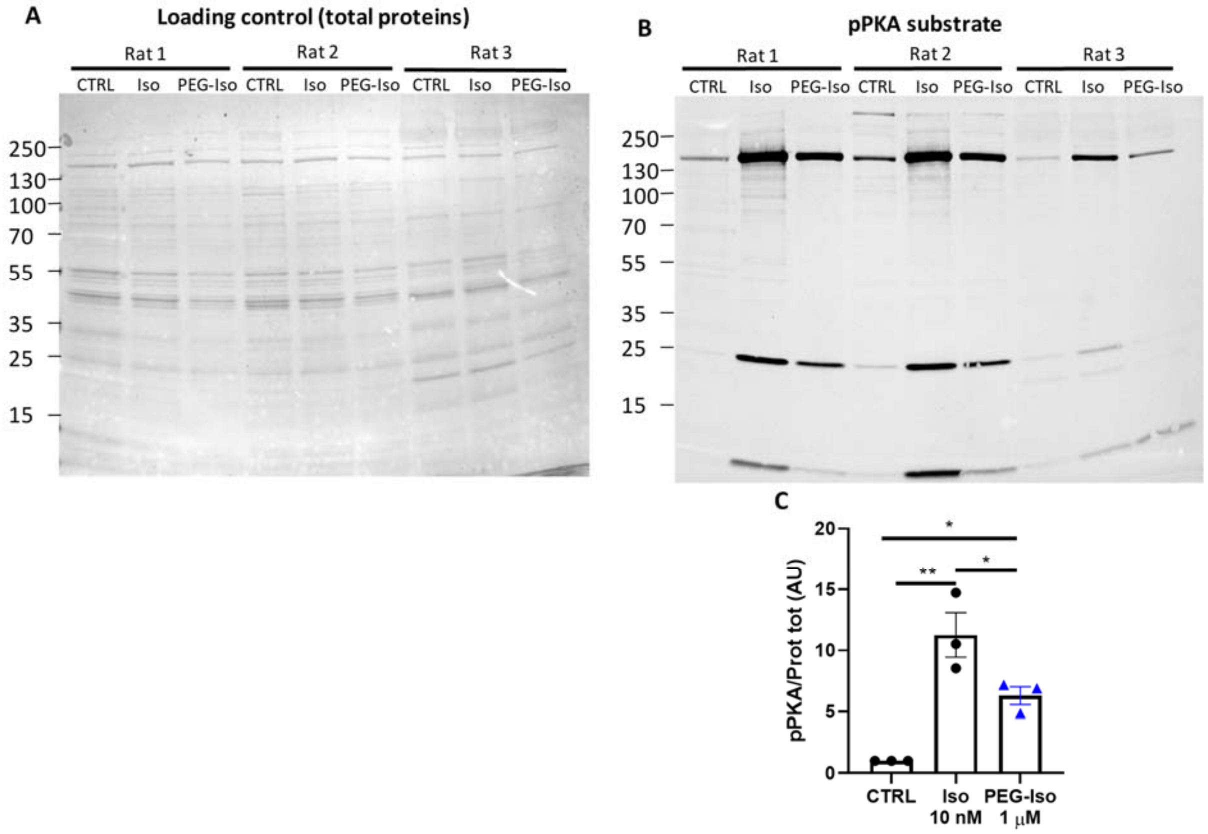


**Fig. S2. Comparison of the effects of PEG-Iso and Iso on nuclear proteins phosphorylation levels by PKA.** Freshly isolated ARVMs were treated with or without Iso 10 nM or PEG-Iso 1 µM during 15 min. Nuclear proteins were extracted, and proteins phosphorylated by PKA were revealed using a phospho-PKA substrate antibody. (**A**) Total proteins labelling as a loading control; (**B**) Immunoblot for phosphorylation of PKA substrates using a specific phospho-PKA antibody; (**C**) Quantification of phospho-PKA levels normalized on total protein, N=3. One-way ANOVA and Tukey's post hoc test: * $p<0.05$; ** $p<0.01$.

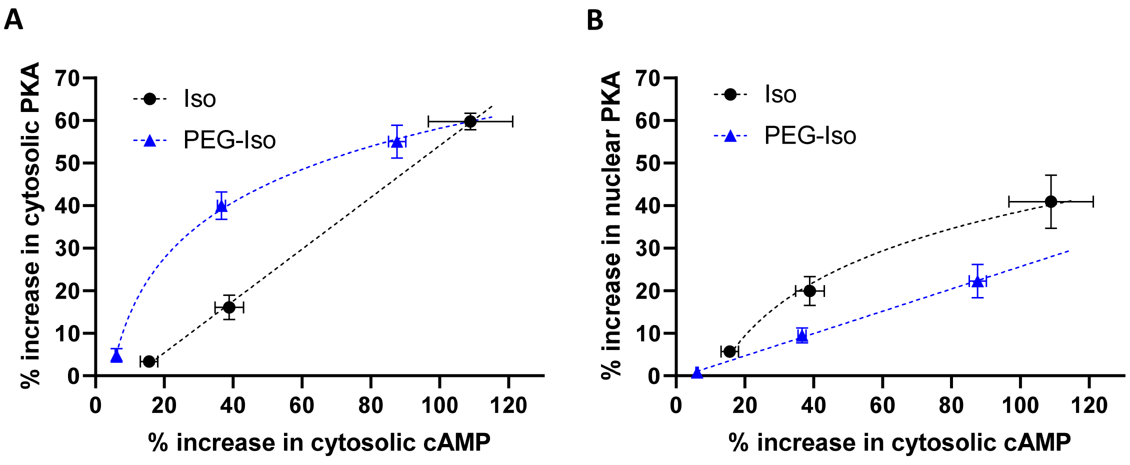


**Fig. S3. Plot of cytosolic and nuclear PKA activity as a function of cytosolic cAMP in response to Iso and PEG-Iso.** For three concentrations of Iso (0.3, 1 and 10 nM) and PEG-Iso (10 nM, 100 nM and 1 µM), the average percent increase (± s.e.m.) in YFP/CFP ratio for cytosolic PKA activity (**A**, taken from Fig. 7) and nuclear PKA activity (**B**, taken from Fig. 8) are plotted as a function of the average percent increase (± s.e.m.) in CFP/YFP ratio for cytosolic cAMP (taken from Fig. 3).
